# Cross-amplified Barcodes on Slides for Spatial Transcriptomics Sequencing

**DOI:** 10.1101/2022.08.25.504658

**Authors:** Zhengyang Jin, Nianzuo Yu, Junda Bai, Zhiyi Liu, Hongrui Li, Junhu Zhang, Chongyang Liang

**Affiliations:** Department of Biopharmacy, The School of Pharmaceutical Sciences, Jilin University, Changchun, 130021, P. R. China; State Key Laboratory of Supramolecular Structure and Materials, College of Chemistry, Jilin University, Changchun, 130012, P. R. China; Joint Laboratory of Opto-Functional Theranostics in Medicine and Chemistry, The First Hospital of Jilin University, Changchun, 130031, P. R. China

**Author notes:** These authors contributed equally to this work (Zhengyang Jin, Nianzuo Yu). To whom correspondence should be addressed. (C.L), (J.Z).

## Abstract

Spatial transcriptomics can reveal molecular signatures of tissue at spatial scales, but current technologies cannot integrate excellent accessibility and easy data-decoding. Here, we use oligonucleotides whose number is the square root of the number of spots to generate cross-amplified spatial barcodes on slides using microfluidic technology. This method can obtain microarrays with well-defined barcodes easily and at low cost, without post-decoding, which contributes to the popularization of spatial transcriptomics.

---

Current spatial transcriptome technologies include in situ RNA sequencing (ISS) based on probe hybridization and image processing, and in situ capture sequencing based on next generation sequencing (NGS)^1^. The former can achieve single molecule level detection at the subcellular level, but the high requirements for equipment, cumbersome experimental and analytical processes, and high costs limit the widespread popularity of such technologies, it also can’t be applied to clinical pathological tissues^2,3^. NGS-based in situ sequencing allows mRNA in tissue sections to be hybridized directly to arrays (e.g., microsphere or microarray) containing spatial barcode primers, allowing spatial location information to be encoded into transcriptomic libraries and sequenced for unbiased transcriptomic detection and visualization of gene spatial expression in tissue sections^4,5^. Among them, microsphere-based methods have achieved subcellular resolution, but the data obtained by such methods are sparse, more importantly, the random distribution of microsphere on slides leads to the need to decode several times to obtain the spatial coordinates, which affects their accessibility and only leads to a laboratory stage^6^. Microarray-based spatial transcriptome technology can obtain more data, but the high-precision automatic micro-spotting facility, which costs millions of dollars, limits its worldwide availability to a wide range of users^7^.

To bridge this gap, we presented cross-amplified barcodes on slides for spatial transcriptomics sequencing (CBSST-Seq), a method to form high-density oligonucleotide microarrays on the surface of slides by cross-amplified reaction without pre-decoding, using standard laboratory instruments and procedures (Fig 1a). Brain tissue was cryosectioned, imaged and permeabilized, and RNA was captured for RNA-seq (Fig 1b), ultimately we demonstrated its application to analyze gene expression at a spatial scale.

**Fig 1.**
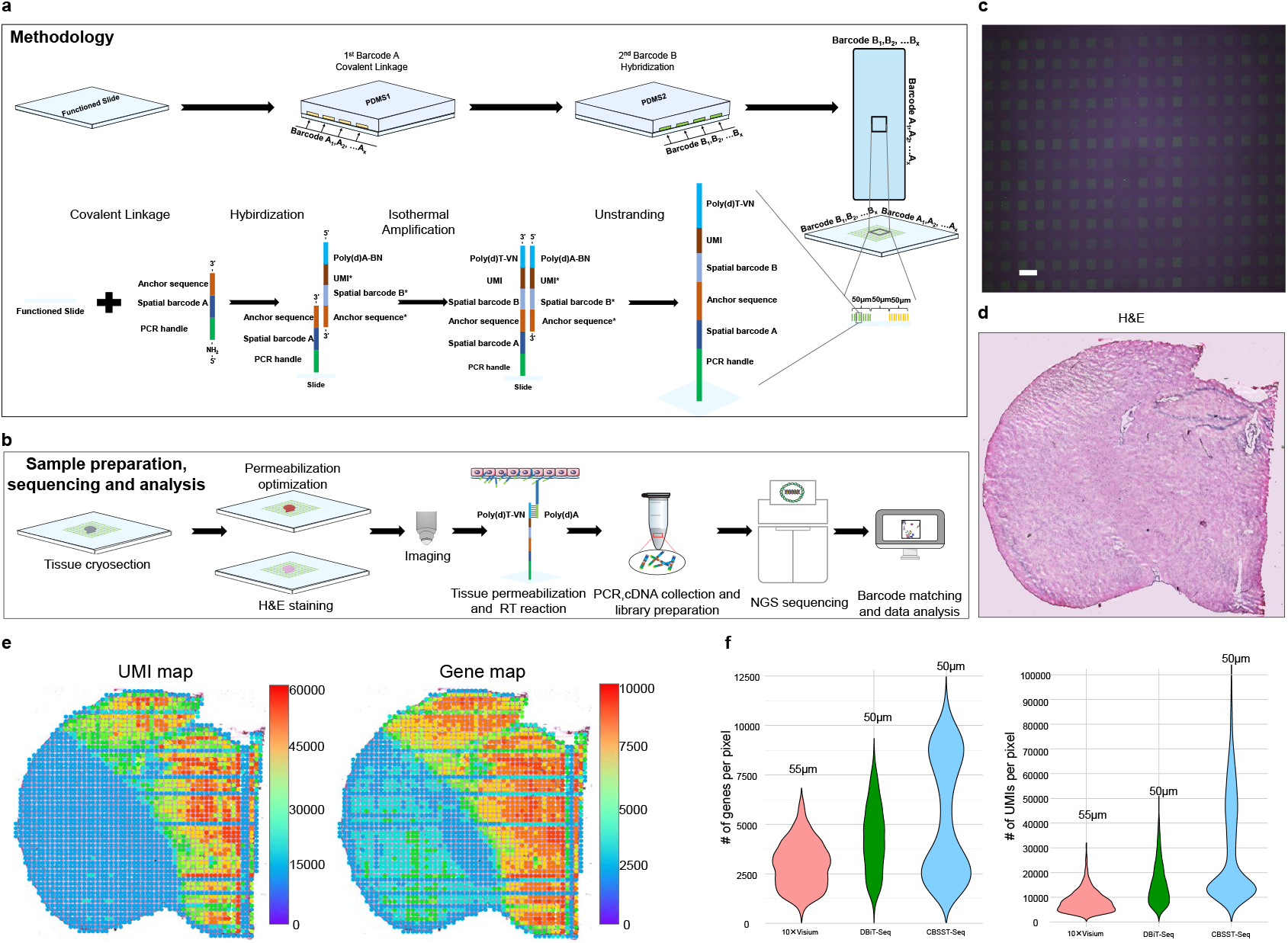
CBSST-Seq. (a) CBSST-Seq Workflow. (b) CBSST-Seq H&E image and Fluorescent footprint image, Scale bar:100μm (c) Validation of spatial barcoding for pixels. The functional slides were spatially barcoded and the pixels characterized by the quality inspection sequence were displayed by fluorescence microscopy (d) The UMI and gene heat maps (pixel size, 50μm), which are closely related to anatomical tissue morphology.(e) Gene and UMI count distribution. CBSST-seq is compared to 10×Visium and DbiT-Seq with the same pixel sizes.

In order to prepare the spatially encoded microarray, microfluidic chip with parallel channels (Supplementary Fig 1a) was placed on the surface of the functional slide, and oligonucleotide sequences containing PCR handle, spatial barcode, anchor sequence and functionalized motifs at the 5’ end for covalent attachment to the slide were introduced. Then we adjusted the channels to be perpendicular to the flow direction of the first set of oligonucleotides and introduced the second set of oligonucleotides containing complementary sequence of anchor sequence, spatial barcode, unique molecular index (UMI) and NB-poly(d) A. This step resulted in the hybridization of the second set of oligonucleotides with the first set of oligonucleotides at the cross-links using the 3 ‘end anchor sequence. By isothermal amplification reaction, microarrays were generated at the intersection of the two sets of barcodes, and the sequence consisted of the 5’ to 3’ ends as: PCR handle, spatial barcode A, anchoring sequence, spatial barcode B, UMI, and poly(d)T-VN (Fig 1a), wherein two sets of known spatial barcodes can refer to the spatial location of the pixel. With this approach, we only need tens of oligonucleotides to generate thousands of pixels, and it does not suffer from surface tension of droplet that limits the resolution and density of microarrays as inkjet printing.

To evaluate the feasibility of the CBSST-Seq workflow, a microfluidic device with a width of 50 μm and a channel count of 70×70: amino-modified oligonucleotides (Supplementary table S1) were introduced into on the surface of N-hydroxysuccinimide slide surface. After the covalent linkage, oligonucleotides with reverse base complementarity to the first set of oligonucleotides were introduced (Supplementary table S1), which are fluorescein-labeled at 5’ ends, and negative control was introduced into the adjacent channels. The results showed that only the channels that produced base reverse complementation could form fluorescent images, and no cross-contamination or fluid leakage were observed (Supplementary Fig 1b). Then we eluted the characterization sequences, switched the channel direction and injected oligonucleotide sequences (Supplementary table S2) that were inversely complementary to the 3’ end anchor sequence of the first set. After incubation, DNA polymerase was added and isothermal amplification reaction was carried out, resulting in pixels where each pixel was a different combination of A and B oligonucleotide sequences. Characterization of pixels was performed using fluorescein-labeled poly(d)A (Supplementary table S2) sequences and it was found that only at the intersection of channels could square fluorescent pixels be generated (Fig 1c).

Validation of microarrays was conducted using frozen sections of mouse brain (Fig 1d). The structural analysis of the PCR amplicons was as expected (Supplementary Fig 1c), and the nucleic acid library size was distributed around 400-700 bp (Supulementary Fig 1d), paired-end dual-indexed sequencing results showed that all different sets of spatial barcodes could be found in the sequencing data. We calculated the total number of UMIs and genes detected per pixel and generated spatial gene map and UMI map (Fig 1e). The results showed that we detected a total of 21324 genes, with an average of 3190 genes captured per pixel point as well as 7133 UMIs, which is comparable to the level of 10×Visium at 55 μm resolution and DBiT-Seq, another spatial transcriptomics technique at 50 μm resolution^7,8^ (Fig 1f).

Further, we generated gene spatial expression patterns of mouse brain tissues and performed cell clustering (Fig 2a, Supplementary Fig 2a). Marker genes such as STX1A in the cortex, Prkcd in the thalamus, HPCA in the hippocampus and Hcrt in the thalamus^9–12^ were visualized (Fig 2b, Supplementary Fig 2b). Based on the H&E staining and markers, the somatic layer and dendritic layer of the hippocampus can be distinguished, such as Prox1 localized in the granule cell layer (DG-sg) of the dentate gyrus, Wfs1 in the CA1 cone layer, and Aldoc in the dendritic layer^13^ (Fig 2c). The strong expression of the astrocytic gene marker Camk2a was also identified in regions within the hippocampus (Supplementary Fig 2c). Meanwhile, the joint expression analysis of several important genes affecting myelin production, such as Cldn11, Plp1 and Mbp showed that they are all located in the dense nerve fiber bundle of myelin sheath^14^ (Fig 2d). We consulted the literature database and the classical Allen Mouse Brain Atlas^15^, as well as marker genes, and performed manual anatomical annotation to reveal the main tissue types (Fig 2e).

**Fig 2.**
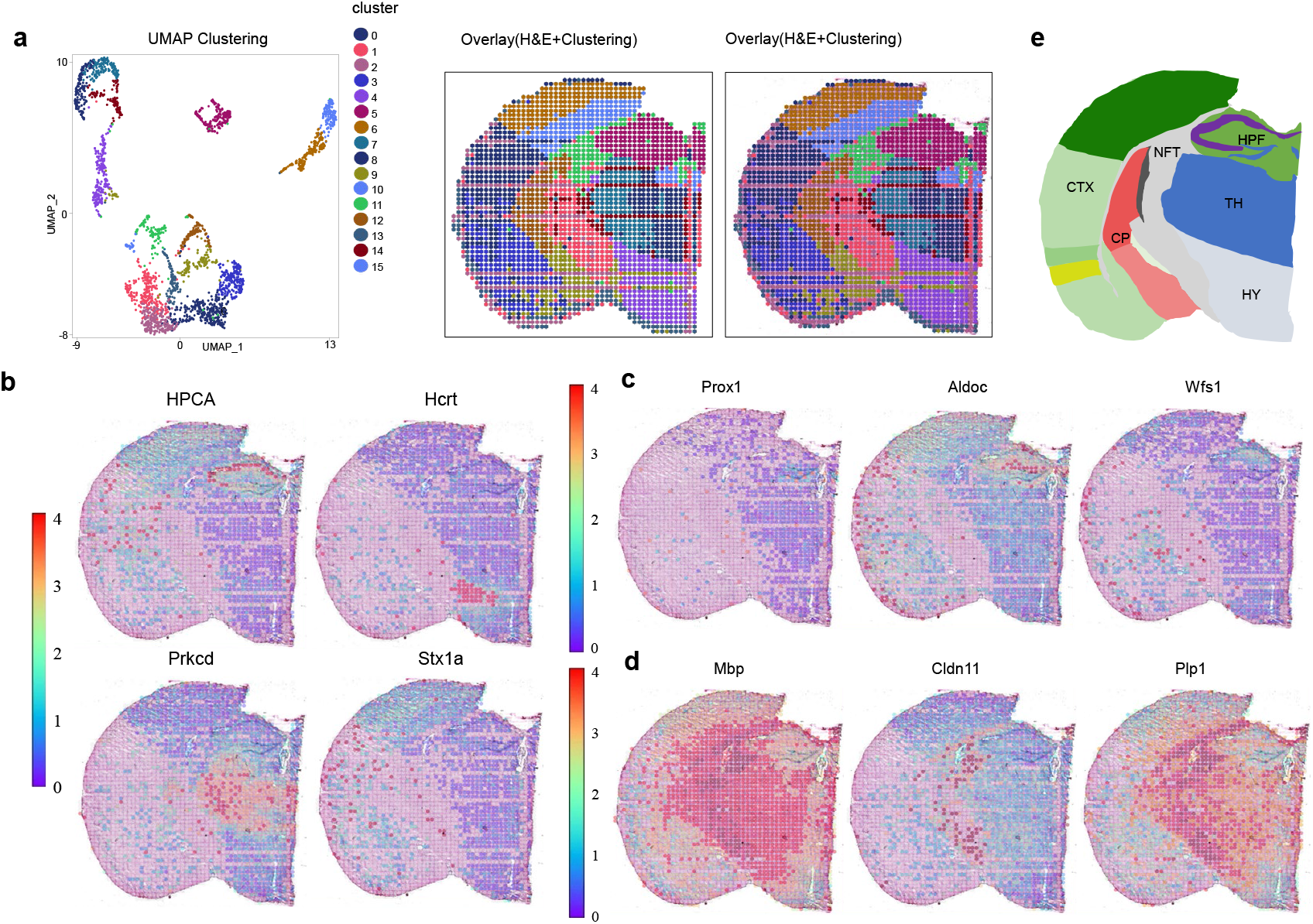
Spatial Gene Expression Mapping of Mouse Brain. (a) Unsupervised clustering analysis and spatial pattern. Left: UMAP showing the clusters of tissue pixel. Middle: spatial distribution of the clusters. Right: overlay of spatial cluster map and tissue image (H&E). (b) Spatial expression of hippocalcin (HPCA), protein kinase C delta (Prkcd), hypocretin neuropeptide precursor (Hcrt) and syntaxin 1A(Stx1a). (c) Spatial expression of prospero homeobox 1 (Prox1) in the granule cell layer of the dentate gyrus, wolframin ER transmembrane glycoprotein (Wfs1) in the CA1 cone layer, and aldolase, fructose-bisphosphate C (Aldoc) in the dendritic layer. (d) The joint expression of claudin 11 (Cldn11), proteolipid protein 1 (Plp1) and myelin basic protein (Mbp). (e) Anatomical annotation of mouse brain. CTX: Cortex, HPF: Hippocampus, CP: Caudate putamen, NFT: Nerve fiber tracts, TH: Thalamus, HY: Hypothalamus.

For additional testing, we analyzed the differentially expressed genes in the hippocampus, thalamus, cortex and hypothalamus, which are anatomically and functionally distinct (Supplementary Fig 3a), even in the cortex of the same anatomical area, there are significant differences between the two adjacent clustering genes. GO and pathway analysis demonstrated the pathways orchestrated by specific genes in different anatomical regions, such as the hippocampus, which is highly associated with chronic depression^16^ (Supplementary Fig 3b,c).

CBSST-Seq substantially improves the accessibility of spatial transcriptome technology while taking into account the resolution, solving one of the drawbacks of spatial transcriptomics, that is, eliminating the need for microprinting technology’s million-dollar startup manufacturing and cumbersome post-decoding and in situ sequencing processes. To facilitate comparison with existing technologies, we do not show the application of CBSST-Seq in higher resolution, but the channel width can be controlled to within 10μm^8^. In conclusion, CBSST-Seq is a simple method that can popularize spatial transcriptome technology and promote rapid multidisciplinary development. It has made a breakthrough in lowering the technical threshold, and it will be conducive to the large-scale popularization of spatial transcriptomics, so as to benefit more researchers.

## Supporting information

CBSST-Seq-SM-Figure

CBSST-Seq-SM-Table

## Acknowledgements

We thank NovelBio for the support of spatial transcriptome sequencing and data analysis.

## Author contributions

ZJ and NY designed the study and wrote the manuscript. ZJ, NY and JB conducted experiments. ZJ, ZL and HL performed bioinformatics analysis.CL and JZ supervised the study and revised the manuscript.CL and JZ are responsible for the overall content as the guarantor.

## Methods

### Microfluidic device fabrication

The glass plate with uniform chromium film and photoresist layer was placed under the mask plate with parallel arrangement of apertures and exposed by UV lamp, and the substrate was soaked in the developing solution to obtain the photoresist glass surface with micro-aperture pattern of spin chromium layer. After that, the surface was immersed in chromium etching solution and placed in glass etching solution (HF, HNO_3_, NH_4_F and H_2_O in the mass ratio of 25:23.5:9.35:450) to obtain a glass aperture mold with the morphological structure of the micro channel pattern. The polydimethylsiloxane (PDMS) prepolymer and the curing agent were mixed in the ratio of 10:1 by mass, degassed under vacuum, poured onto the surface of the glass mold, and cured in an oven at a temperature of 60°C. Finally, the PDMS microfluidic pore channels arranged in parallel were uncovered, and the width of the pore channels and the spacing between the pore channels were measured precisely.

### Preparation of microfluidic-based spatially encoded slides

We used two sets of oligonucleotides to generate microarrays. Wherein the first set of oligonucleotides comtaining: (1) PCR handle,(2) a 9 base pair(bp)’Spatial barcode A’,(3) a 12bp anchor sequence(/AmC6/CTACACGACGCTCTTCCGA-Spatial barcode A-ACTGGCCTGCGA) (Genscript). To enlarge the barcode pool, we introduced the second set of oligonucleotides including: (1) a 20bp NB-poly(d)A,(2) a 8bp UMI,(3) a 9bp reverse complementary sequence of ‘Spatial barcode B’ and(4) a 12bp reverse complementary sequence of anchor sequence (NBAAAAAAAAAAAAAAAAAA-NNNNNNNN-Spatial barcode B*-TCGCAGGCCAGT). The parallel arranged microchannels were attached to the surface of slides(PolyAn) coated with N-hydroxysuccinimide(NHS)groups, then the first set of oligonucleotides(50μM) were introduced into the channels and incubated in a wet box overnight to covalently connect them with the slide surface. The washing buffer (PBS,pH=7.4) was introduced to wash away the excess nucleic acid and dry the slide. The microchannels were reapplied to the slide surface in an orientation perpendicular to the above and the second set of oligonucleotides(10μM) were introduced, which can be complementarily hybridized to the first set of oligonucleotides by the reverse complementary sequence of the anchoring sequence. An isothermal extension reaction mixture was made from 35μl NEBuffer2 (10×, New England Biolabs), 12.5μl Klenow Fragment (5,000U/ml, New England Biolabs), 7μl of dNTPs (10mM, New England Biolabs), 295.5μl Nuclease-Free Water (Thermo Fisher). The mix was then introduced into the microchannels and incubated at 37°C for half an hour to generate the oligonucleotide microarray with spatial coordinates. The nucleic acid sequences of the microarray consist of the 5’ to 3’ ends as: PCR handle, spatial barcode A, anchor sequence, spatial barcode B, UMI, poly(d)T-VN (/AmC6/ CTACACGACGCTCTTCCGA-Spatial barcode A–ACTGGCCTGCGA-Spatial barcode B– NNNNNNNNTTTTTTTTTTTTTTTTTTVN). The reaction was terminated by incubation with 10 mM EDTA for 10 min at 75°C. The reaction mixture was washed off and the melting mixture (150mM NaOH) was added on the microarray surface to remove the second group of oligonucleotide sequences.

### Quality assessment of spatially coded slides

After connecting the first set of oligonucleotide sequences on the surface of the slide, we used the reverse complementary sequence of anchor sequence with 5’end FAM as a positive control (10μM), and introduced the negative control sequence into the channel alternately. After half an hour of incubation, PBS was used for cleaning, and then fluorescence microscopy (Olympus) was used for imaging to characterize the linkage of of the first set of oligonucleotides and the leakage of the microchannels. After generating the microarray, the 5’ end fluorescein-labeled poly(d)A sequence was added dropwise on the slide surface, incubated for half an hour to characterize the generation of oligonucleotide sequence microarray using fluorescence microscopy.

### Tissue samples

We followed a protocol as described in Ståhl et al^1^. Adult C57BL/6J mice (at 10 weeks of age) were euthanized and their brain tissue dissected. It was then frozen in an isopentane bath at −40°C and transferred to −80°C for storage. Frozen tissues were embedded in pre-cooled OCT (Sakura) and frozen sections were taken at 10 μm thickness.

This study complied with all relevant ethical regulations regarding experiments on animal tissue samples.

### Tissue staining and imaging

The tissue sample was attached to the surface of the slide, and then placed at 37°C for 1min, and then transferred into the precooled methanol for 30min at −20°C. Sections were incubated for 5min with hematoxylin (SigmaAldrich) followed by 2 min in bluing buffer (DAKO) and 10s in eosin Y (0.45M, pH6). The images were taken via the automatic digital slide scanner Pannoramic MIDI (3DHISTECH) at × 20 resolution.

### Spatial transcriptomics

We followed the protocol of 10xGenomics for tissue optimization experiments (10xGenomics, Visium Spatial Tissue Optimization, CG000238 Rev A). Fluorescence footprint imaging was performed using the automatic digital slide scanner to optimize the permeabilization conditions. Then, we performed spatial transcriptome experiment on the tissue samples following the standard procedure of Visum Spatial Gene Expression Reagent Kits (10x genomics, CG000239, Rev D). The H&E stained slides were permeabilized with tissue permeabilizing enzyme (12min), then incubated with RT master mix for 45 min, after washing and then incubated with the second cha in mixture at 65°C for 15min. 0.1×SSC was used for cleaning between steps, and KOH was used for collecting the second chain. qPCR was performed using KAPA SYBR FAST qPCR Master Mix (KAPA Biosystems) and analyzed using QuantStudio 6 Flex (Thermo Fisher). The Cq value of 25% of the peak fluorescence was used as the cycle number of cDNA amplification. DNA fragmentation, end repair, A-tailing and adaptor ligation were performed according to the instructions. Libraries were constructed using the Kapa HIFI HotStart ReadyMix PCR Kit (KAPA Biosystems). Nucleic acid purification was performed using SPRI Select (BECKMAN COULTER), nucleic acid concentration was determined by Qubit High Sensitivity DNA assay (Thermo Fisher), and fragment length analysis was performed using 2200 TapeStation (Agilent Technologies). The libraries were finally sequenced using an Illumina Novaseq6000 sequencer and 150 pair-end double index setting was used for sequencing.

### CBSST-Seq raw data processing

Raw data was processed using fastp, filtering adaptor sequences with default parameters and removing low quality reads^2^.

### Data analysis

The filtered data was mapped to the mouse genome (mm10 mouse) using SpaceRanger V1.1.0 to obtain UMI counts, feature-barcode matrix and determine clusters for down-stream analysis. Raw UMI count spot matrices, images, spot-image coordinates and scale factors were imported into R 4.2.1. Pixel locations were determined by spatial barcodes A and B. Spots overlaying tissue sections were kept by filtering out the spot matrix. Heatmaps were generated using unnormalized feature and UMI counts. Graphcluster was utilized for spatial spot clustering and Wilcox rank sum test was used for marker gene analysis. Pixels containing a total number of detected genes < 200, and those with a total number of detected genes ranking in the top 0.1% were filtered out. The Seurat V3.2 package was applied to normalization and sctransform, as recommended in Seurat package.PCA was used for dimensionality reduction based on 2,000 highly variable genes. Unsupervised cell clusters were acquired by graph-based clustering approach (the top 30 PCs were selected, resolution = 0.5). Visualization was realized by T-SNE and UMAP. Tissue regions were annotated based on H&E stained brightfield images and marker genes. For marker gene expression, we used Seurat and Wilcoxon rank sum test for statistical testing^3^. FindAllMarkers was used to identify differentially expressed gene(logFC>0.25; min.pct>0.25; adjusted P value < 0.05.). SpatialDE was used to identify spatially distributed genes, and results were further used for GO and pathway analysis^4^.

### Comparison of CBSST-Seq with published methods

DBiT-Seq data were taken from GSE137986^5^, and Visium data from GSE153859^6^.

### Data availability

The accession number for the sequencing data reported in this paper is submitted to GEO: https://www.ncbi.nlm.nih.gov/geo/query/acc.cgi?acc=GSE211858.

