## Supplementary material for "Cross-amplified Barcodes on Slides for Spatial Transcriptomics Sequencing": CBSST-Seq-SM-Figure

**Supplementary Table.S1**

| A1-1 | /AMC6/-CTACACGACGCTCTTCCGATCTCTTCACATAACTGGCCTGCGA |
| --- | --- |
| A1-2 | /AMC6/-CTACACGACGCTCTTCCGATCTACACGCCGGACTGGCCTGCGA |
| A1-3 | /AMC6/-CTACACGACGCTCTTCCGATCTCGGTCCAGGACTGGCCTGCGA |
| A1-4 | /AMC6/-CTACACGACGCTCTTCCGATCTAATCGAATGACTGGCCTGCGA |
| A1-5 | /AMC6/-CTACACGACGCTCTTCCGATCTCCTAGTATAACTGGCCTGCGA |
| A1-6 | /AMC6/-CTACACGACGCTCTTCCGATCTATTGGCTAAACTGGCCTGCGA |
| A1-7 | /AMC6/-CTACACGACGCTCTTCCGATCTAAGACATGCACTGGCCTGCGA |
| A1-8 | /AMC6/-CTACACGACGCTCTTCCGATCTAAGGCGATCACTGGCCTGCGA |
| A1-9 | /AMC6/-CTACACGACGCTCTTCCGATCTGTGTCCTTAACTGGCCTGCGA |
| A1-10 | /AMC6/-CTACACGACGCTCTTCCGATCTGGATTAGGAACTGGCCTGCGA |
| A1-11 | /AMC6/-CTACACGACGCTCTTCCGATCTATGGATCCAACTGGCCTGCGA |
| A1-12 | /AMC6/-CTACACGACGCTCTTCCGATCTACATAAGCGACTGGCCTGCGA |
| A1-13 | /AMC6/-CTACACGACGCTCTTCCGATCTAACTGTATTACTGGCCTGCGA |
| A1-14 | /AMC6/-CTACACGACGCTCTTCCGATCTACCTTGCGGACTGGCCTGCGA |
| A1-15 | /AMC6/-CTACACGACGCTCTTCCGATCTCAGGTGTAGACTGGCCTGCGA |
| A1-16 | /AMC6/-CTACACGACGCTCTTCCGATCTAGGAGATTAACTGGCCTGCGA |
| A1-17 | /AMC6/-CTACACGACGCTCTTCCGATCTGCGATTACAACTGGCCTGCGA |
| A1-18 | /AMC6/-CTACACGACGCTCTTCCGATCTACCGGATAGACTGGCCTGCGA |
| A1-19 | /AMC6/-CTACACGACGCTCTTCCGATCTCCACTTGGAACTGGCCTGCGA |
| A1-20 | /AMC6/-CTACACGACGCTCTTCCGATCTAGAGAAGTTACTGGCCTGCGA |
| A1-21 | /AMC6/-CTACACGACGCTCTTCCGATCTTAAGTTCGAACTGGCCTGCGA |
| A1-22 | /AMC6/-CTACACGACGCTCTTCCGATCTACGGATATTACTGGCCTGCGA |
| A1-23 | /AMC6/-CTACACGACGCTCTTCCGATCTTGGCTCAGAACTGGCCTGCGA |
| A1-24 | /AMC6/-CTACACGACGCTCTTCCGATCTGAATCTGTAACTGGCCTGCGA |
| A1-25 | /AMC6/-CTACACGACGCTCTTCCGATCTACCAAGGACACTGGCCTGCGA |
| A1-26 | /AMC6/-CTACACGACGCTCTTCCGATCTAGTATCTGTACTGGCCTGCGA |
| A1-27 | /AMC6/-CTACACGACGCTCTTCCGATCTCACACACTAACTGGCCTGCGA |
| A1-28 | /AMC6/-CTACACGACGCTCTTCCGATCTATTAAGTGCACTGGCCTGCGA |
| A1-29 | /AMC6/-CTACACGACGCTCTTCCGATCTAAGTAACCCACTGGCCTGCGA |
| A1-30 | /AMC6/-CTACACGACGCTCTTCCGATCTAAATCCTGTACTGGCCTGCGA |
| A1-31 | /AMC6/-CTACACGACGCTCTTCCGATCTCACATTGCAACTGGCCTGCGA |
| A1-32 | /AMC6/-CTACACGACGCTCTTCCGATCTGCACTGTCAACTGGCCTGCGA |
| A1-33 | /AMC6/-CTACACGACGCTCTTCCGATCTATACTTAGGACTGGCCTGCGA |
| A1-34 | /AMC6/-CTACACGACGCTCTTCCGATCTGCAATCCGAACTGGCCTGCGA |
| A1-35 | /AMC6/-CTACACGACGCTCTTCCGATCTACGCAATCAACTGGCCTGCGA |
| A1-36 | /AMC6/-CTACACGACGCTCTTCCGATCTGAGTATTAGACTGGCCTGCGA |
| A1-37 | /AMC6/-CTACACGACGCTCTTCCGATCTGACGGATTAACTGGCCTGCGA |
| A1-38 | /AMC6/-CTACACGACGCTCTTCCGATCTCAGCTGACAACTGGCCTGCGA |
| A1-39 | /AMC6/-CTACACGACGCTCTTCCGATCTCAACATATTACTGGCCTGCGA |
| A1-40 | /AMC6/-CTACACGACGCTCTTCCGATCTAACTTCTCCACTGGCCTGCGA |
| A1-41 | /AMC6/-CTACACGACGCTCTTCCGATCTCTATGAAATACTGGCCTGCGA |
| A1-42 | /AMC6/-CTACACGACGCTCTTCCGATCTATTATTACCACTGGCCTGCGA |
| A1-43 | /AMC6/-CTACACGACGCTCTTCCGATCTTACCGAGCAACTGGCCTGCGA |
| A1-44 | /AMC6/-CTACACGACGCTCTTCCGATCTTCTCTTCAAACTGGCCTGCGA |
| A1-45 | /AMC6/-CTACACGACGCTCTTCCGATCTTAAGCGTTAACTGGCCTGCGA |
| A1-46 | /AMC6/-CTACACGACGCTCTTCCGATCTGCCTTACAAACTGGCCTGCGA |
| A1-47 | /AMC6/-CTACACGACGCTCTTCCGATCTAGCACACAGACTGGCCTGCGA |
| A1-48 | /AMC6/-CTACACGACGCTCTTCCGATCTACAGTTCCGACTGGCCTGCGA |
| A1-49 | /AMC6/-CTACACGACGCTCTTCCGATCTAGTAAAGCCACTGGCCTGCGA |
| A1-50 | /AMC6/-CTACACGACGCTCTTCCGATCTCAGTTTCACACTGGCCTGCGA |
| A1-51 | /AMC6/-CTACACGACGCTCTTCCGATCTCGTTACTAAACTGGCCTGCGA |
| A1-52 | /AMC6/-CTACACGACGCTCTTCCGATCTTTGTTCCAAACTGGCCTGCGA |
| A1-53 | /AMC6/-CTACACGACGCTCTTCCGATCTAGAAGCACTACTGGCCTGCGA |
| A1-54 | /AMC6/-CTACACGACGCTCTTCCGATCTCAGCAAGATACTGGCCTGCGA |
| A1-55 | /AMC6/-CTACACGACGCTCTTCCGATCTCAAACCGCCACTGGCCTGCGA |
| A1-56 | /AMC6/-CTACACGACGCTCTTCCGATCTCTAACTCGCACTGGCCTGCGA |
| A1-57 | /AMC6/-CTACACGACGCTCTTCCGATCTAATATTGGGACTGGCCTGCGA |
| A1-58 | /AMC6/-CTACACGACGCTCTTCCGATCTAGAACTTCCACTGGCCTGCGA |
| A1-59 | /AMC6/-CTACACGACGCTCTTCCGATCTCAAAGGCACACTGGCCTGCGA |
| A1-60 | /AMC6/-CTACACGACGCTCTTCCGATCTAAGCTCAACACTGGCCTGCGA |
| A1-61 | /AMC6/-CTACACGACGCTCTTCCGATCTTCCAGTCGAACTGGCCTGCGA |
| A1-62 | /AMC6/-CTACACGACGCTCTTCCGATCTAGCCATCACACTGGCCTGCGA |
| A1-63 | /AMC6/-CTACACGACGCTCTTCCGATCTAACGAGAAGACTGGCCTGCGA |
| A1-64 | /AMC6/-CTACACGACGCTCTTCCGATCTCTACAGAACACTGGCCTGCGA |
| A1-65 | /AMC6/-CTACACGACGCTCTTCCGATCTAGAGCTATGACTGGCCTGCGA |
| A1-66 | /AMC6/-CTACACGACGCTCTTCCGATCTGAGGATGGAACTGGCCTGCGA |
| A1-67 | /AMC6/-CTACACGACGCTCTTCCGATCTTGTACCTTAACTGGCCTGCGA |
| A1-68 | /AMC6/-CTACACGACGCTCTTCCGATCTACACACAAAACTGGCCTGCGA |
| A1-69 | /AMC6/-CTACACGACGCTCTTCCGATCTTCAGGAGGAACTGGCCTGCGA |
| A1-70 | /AMC6/-CTACACGACGCTCTTCCGATCTGAGGTGCTAACTGGCCTGCGA |
| A-Control | FAM-TCGCAGGCCAGTTATGTGAAG |

**Talbe S1.** The first group of oligonucleotide sequences and characterization sequences for CBSST-Seq.

**Supplementary Table.S2**

| B2-1 | NBAAAAAAAAAAAAAAAAAANNNNNNNNATACTTGTGTCGCAGGCCAGT |
| --- | --- |
| B2-2 | NBAAAAAAAAAAAAAAAAAANNNNNNNNCTAAGGTGTTCGCAGGCCAGT |
| B2-3 | NBAAAAAAAAAAAAAAAAAANNNNNNNNTTGTCGTTCTCGCAGGCCAGT |
| B2-4 | NBAAAAAAAAAAAAAAAAAANNNNNNNNGTACAGACTTCGCAGGCCAGT |
| B2-5 | NBAAAAAAAAAAAAAAAAAANNNNNNNNCTGTAATTTTCGCAGGCCAGT |
| B2-6 | NBAAAAAAAAAAAAAAAAAANNNNNNNNTCTGTAGCCTCGCAGGCCAGT |
| B2-7 | NBAAAAAAAAAAAAAAAAAANNNNNNNNCGATACATTTCGCAGGCCAGT |
| B2-8 | NBAAAAAAAAAAAAAAAAAANNNNNNNNTTCTACTTGTCGCAGGCCAGT |
| B2-9 | NBAAAAAAAAAAAAAAAAAANNNNNNNNTAAGAGATCTCGCAGGCCAGT |
| B2-10 | NBAAAAAAAAAAAAAAAAAANNNNNNNNCGCGTTGTTTCGCAGGCCAGT |
| B2-11 | NBAAAAAAAAAAAAAAAAAANNNNNNNNTAACTCACCTCGCAGGCCAGT |
| B2-12 | NBAAAAAAAAAAAAAAAAAANNNNNNNNCCCTCCCTGTCGCAGGCCAGT |
| B2-13 | NBAAAAAAAAAAAAAAAAAANNNNNNNNTAAGACGGATCGCAGGCCAGT |
| B2-14 | NBAAAAAAAAAAAAAAAAAANNNNNNNNTACTATGCATCGCAGGCCAGT |
| B2-15 | NBAAAAAAAAAAAAAAAAAANNNNNNNNATCGTAAGTTCGCAGGCCAGT |
| B2-16 | NBAAAAAAAAAAAAAAAAAANNNNNNNNTCGCATACATCGCAGGCCAGT |
| B2-17 | NBAAAAAAAAAAAAAAAAAANNNNNNNNTCAAGGAGCTCGCAGGCCAGT |
| B2-18 | NBAAAAAAAAAAAAAAAAAANNNNNNNNTGTTGTGCCTCGCAGGCCAGT |
| B2-19 | NBAAAAAAAAAAAAAAAAAANNNNNNNNTGTCTTGAGTCGCAGGCCAGT |
| B2-20 | NBAAAAAAAAAAAAAAAAAANNNNNNNNCAACAGCGTTCGCAGGCCAGT |
| B2-21 | NBAAAAAAAAAAAAAAAAAANNNNNNNNTTACAATATTCGCAGGCCAGT |
| B2-22 | NBAAAAAAAAAAAAAAAAAANNNNNNNNCGTAAACTTTCGCAGGCCAGT |
| B2-23 | NBAAAAAAAAAAAAAAAAAANNNNNNNNGCCAGGCTGTCGCAGGCCAGT |
| B2-24 | NBAAAAAAAAAAAAAAAAAANNNNNNNNGGCTAATAGTCGCAGGCCAGT |
| B2-25 | NBAAAAAAAAAAAAAAAAAANNNNNNNNCCACGTTTGTCGCAGGCCAGT |
| B2-26 | NBAAAAAAAAAAAAAAAAAANNNNNNNNAATGACTTTTCGCAGGCCAGT |
| B2-27 | NBAAAAAAAAAAAAAAAAAANNNNNNNNTGCCAAGACTCGCAGGCCAGT |
| B2-28 | NBAAAAAAAAAAAAAAAAAANNNNNNNNTCGCTGATCTCGCAGGCCAGT |
| B2-29 | NBAAAAAAAAAAAAAAAAAANNNNNNNNGCCGAATGTTCGCAGGCCAGT |
| B2-30 | NBAAAAAAAAAAAAAAAAAANNNNNNNNCTAATTACTTCGCAGGCCAGT |
| B2-31 | NBAAAAAAAAAAAAAAAAAANNNNNNNNTTGGCTTCATCGCAGGCCAGT |
| B2-32 | NBAAAAAAAAAAAAAAAAAANNNNNNNNTGTCGTAGATCGCAGGCCAGT |
| B2-33 | NBAAAAAAAAAAAAAAAAAANNNNNNNNAACGTTATGTCGCAGGCCAGT |
| B2-34 | NBAAAAAAAAAAAAAAAAAANNNNNNNNGAGTCCCATTCGCAGGCCAGT |
| B2-35 | NBAAAAAAAAAAAAAAAAAANNNNNNNNTCCTCTATCTCGCAGGCCAGT |
| B2-36 | NBAAAAAAAAAAAAAAAAAANNNNNNNNCGCATGTAGTCGCAGGCCAGT |
| B2-37 | NBAAAAAAAAAAAAAAAAAANNNNNNNNAGATCGTTGTCGCAGGCCAGT |
| B2-38 | NBAAAAAAAAAAAAAAAAAANNNNNNNNTAGGCTAACTCGCAGGCCAGT |
| B2-39 | NBAAAAAAAAAAAAAAAAAANNNNNNNNGATGCAACTTCGCAGGCCAGT |
| B2-40 | NBAAAAAAAAAAAAAAAAAANNNNNNNNAGTTCCCTTTCGCAGGCCAGT |
| B2-41 | NBAAAAAAAAAAAAAAAAAANNNNNNNNATATGTAGTTCGCAGGCCAGT |
| B2-42 | NBAAAAAAAAAAAAAAAAAANNNNNNNNGAAGCTTAGTCGCAGGCCAGT |
| B2-43 | NBAAAAAAAAAAAAAAAAAANNNNNNNNCTGGTTCGTTCGCAGGCCAGT |
| B2-44 | NBAAAAAAAAAAAAAAAAAANNNNNNNNTCCGAAGTATCGCAGGCCAGT |
| B2-45 | NBAAAAAAAAAAAAAAAAAANNNNNNNNATGGATGTTTCGCAGGCCAGT |
| B2-46 | NBAAAAAAAAAAAAAAAAAANNNNNNNNAACCAGGCTTCGCAGGCCAGT |
| B2-47 | NBAAAAAAAAAAAAAAAAAANNNNNNNNGGAAACTTGTCGCAGGCCAGT |
| B2-48 | NBAAAAAAAAAAAAAAAAAANNNNNNNNAAATGCCTGTCGCAGGCCAGT |
| B2-49 | NBAAAAAAAAAAAAAAAAAANNNNNNNNCTCCCACGTTCGCAGGCCAGT |
| B2-50 | NBAAAAAAAAAAAAAAAAAANNNNNNNNTCCGTGAGATCGCAGGCCAGT |
| B2-51 | NBAAAAAAAAAAAAAAAAAANNNNNNNNTAATGTTGCTCGCAGGCCAGT |
| B2-52 | NBAAAAAAAAAAAAAAAAAANNNNNNNNACGGACCATTCGCAGGCCAGT |
| B2-53 | NBAAAAAAAAAAAAAAAAAANNNNNNNNTCATGATAGTCGCAGGCCAGT |
| B2-54 | NBAAAAAAAAAAAAAAAAAANNNNNNNNCTTGTATTGTCGCAGGCCAGT |
| B2-55 | NBAAAAAAAAAAAAAAAAAANNNNNNNNGGCCTCTTTTCGCAGGCCAGT |
| B2-56 | NBAAAAAAAAAAAAAAAAAANNNNNNNNTGCTTCTACTCGCAGGCCAGT |
| B2-57 | NBAAAAAAAAAAAAAAAAAANNNNNNNNTTCCATAGCTCGCAGGCCAGT |
| B2-58 | NBAAAAAAAAAAAAAAAAAANNNNNNNNCCCTGGAGTTCGCAGGCCAGT |
| B2-59 | NBAAAAAAAAAAAAAAAAAANNNNNNNNTGCACTTGTTCGCAGGCCAGT |
| B2-60 | NBAAAAAAAAAAAAAAAAAANNNNNNNNTGGACCATCTCGCAGGCCAGT |
| B2-61 | NBAAAAAAAAAAAAAAAAAANNNNNNNNTATTGAGGATCGCAGGCCAGT |
| B2-62 | NBAAAAAAAAAAAAAAAAAANNNNNNNNTTGTTTATTTCGCAGGCCAGT |
| B2-63 | NBAAAAAAAAAAAAAAAAAANNNNNNNNTCCGTACAGTCGCAGGCCAGT |
| B2-64 | NBAAAAAAAAAAAAAAAAAANNNNNNNNTCTATCTAGTCGCAGGCCAGT |
| B2-65 | NBAAAAAAAAAAAAAAAAAANNNNNNNNCACATAGCTTCGCAGGCCAGT |
| B2-66 | NBAAAAAAAAAAAAAAAAAANNNNNNNNCCTCCATTTTCGCAGGCCAGT |
| B2-67 | NBAAAAAAAAAAAAAAAAAANNNNNNNNCTTGCGGCTTCGCAGGCCAGT |
| B2-68 | NBAAAAAAAAAAAAAAAAAANNNNNNNNGTTTACTGTTCGCAGGCCAGT |
| B2-69 | NBAAAAAAAAAAAAAAAAAANNNNNNNNTCACACGTTTCGCAGGCCAGT |
| B2-70 | NBAAAAAAAAAAAAAAAAAANNNNNNNNGAATTCAGTTCGCAGGCCAGT |
| B-Control | FAM-CGGCTAAAAAAAAAAAAAAAAAA |

**Talbe S2.** The second group of oligonucleotide sequences and characterization sequences for CBSST-Seq.
