## Supplementary material for "Cross-amplified Barcodes on Slides for Spatial Transcriptomics Sequencing": CBSST-Seq-SM-Table

**Supplementary Figure .1**

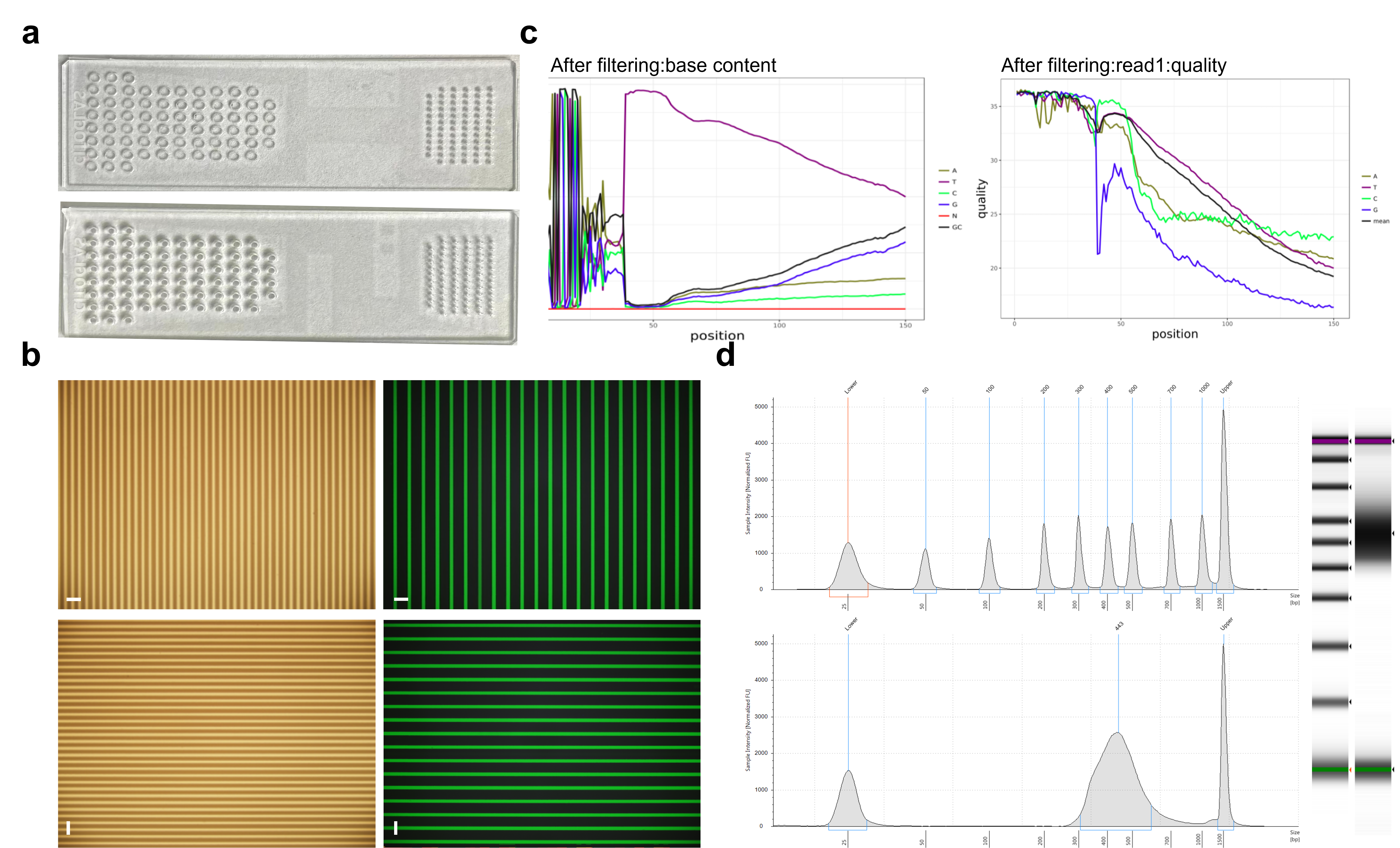

**Figure S1. CBSST-Seq quality** **characterization.** (a) Microfluidic device. (b) Characterization of oligonucleotides using fam labeled fluorescent probes.Scale bar:200μm. (c) Characterization of sequence structure and quality.(D) Length analysis of nucleic acid library.

**Supplementary Figure .2**

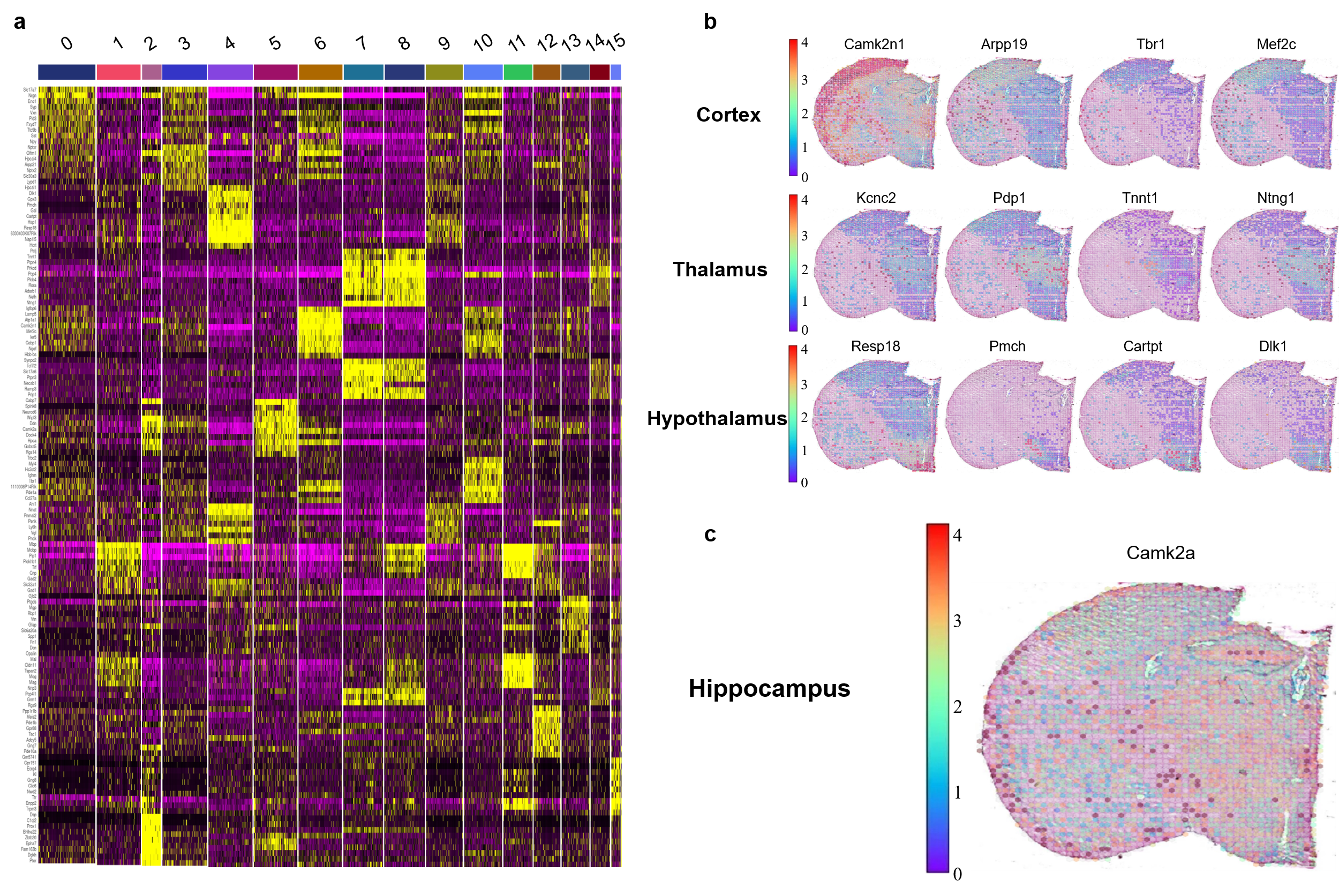

**Figure S2. CBSST-Seq identifies different spatial clusters of mouse brain and gene expression patterns.** (a) Gene expression heatmap of 16 clusters obtained by unsupervised clustering analysis. Top ranked differentially expressed genes are shown in each cluster. (b) Marker genes located in cerebral cortex, thalamus and hypothalamus. (c) Characterization of calcium/calmodulin dependent protein kinase II alpha (Camk2a) gene expression.

**Supplementary Figure .3**

**Figure S3.** **Differential gene analysis, go and pathway analysis of specific anatomical regions of mouse brain.** (a) Differential gene analysis analysis between different anatomical regions. (b)GO analysis analysis between different anatomical regions. (c) Pathway analysis analysis between different anatomical regions.
